# Molecular Heterosis in the Abaca (*Musa textilis* Née) BC_2_ Hybrid, Bandala

**DOI:** 10.1101/2025.04.21.649768

**Authors:** Nelzo C. Ereful, Roneil Christian S. Alonday, Antonio G. Lalusin

## Abstract

The Philippines supplies ∼85% of the global demand for abaca (*Musa textilis* Née) fiber. A backcross (BC_2_) hybrid, named Bandala, was created by crossing Abuab, an abaca variety with high fiber quality, to Pacol, a wild banana (*Musa balbisiana* Colla) variety with resistance against Abaca Bunchy Top Virus (ABTV). To assess regulatory difference and expression heterosis, the parents (Abuab and Pacol) and their BC_2_ were sequenced using RNA-seq. Analysis of expression heterosis showed that a large number of transcripts exhibited non-additive (dominance and transgressive) mode, accounting for ∼83.2% of the total heterotic genes. High-parent Abuab dominant genes in the BC_2_ were identified including genes encoding for cellulose synthase. Results indicated that the combined trans and cis + trans (synergistic interaction) largely explain the combined evolutionary divergence between the parents and repeated backcrossing–selection procedures. Genes exhibiting compensatory interaction are significantly enriched under the transgressive mode of inheritance, contributing largely to the heterotic effect in the backcross under directional selection. Further statistical analysis suggests that regulatory differences strongly influence expression heterosis. We further fitted fixed line equations describing each mode of expression inheritance in a 3D plot. In our concurrent work, the BC_2_ demonstrated overdominance over its parents on two economically important phenotypic traits – fiber length and tensile strength. This study demonstrated that phenotypic and expression heteroses are inherent even in backcrosses owing to the presence of the two alleles in the Bandala genome, providing insights into the genetic mechanisms underlying the heterotic performance of BC_2_ and offering valuable directions for abaca breeding.

**Key Message:** A backcross hybrid (BC_2_) derived by crossing *Musa textilis* var. Abuab and *M. balbisiana* var. Pacol exhibited expression heteroses which can be explained by the underlying cis and/or trans regulatory factors and their interactions.

## Introduction

The Philippines is the leading exporter of fiber derived from abaca (*M. textilis* Née, T genome), also known as Manila hemp, with around 85% of the global demand being supplied by the country. Its demand is expected to further increase across various sectors including maritime, banking, healthcare, automotive, among others, because of its superior mechanical properties.

Abaca, which belongs to the family Musaceae, is host to several pathogens including abaca bunchy top (ABTV), bract mosaic (ABMV), and mosaic viruses (AMV), the three most important disease viruses causing reduction in fiber yield (Halos 2008). As such, there is a need to improve the crop to make it resistant against pathogenic diseases. To partially address this issue, backcrosses derived from Abuab, an abaca variety, were developed to introgress virus-resistant features into its genome.

There has been an increasing number of research investigations accorded to the crop in recent years owing to the recent developments on sequencing technology. For example, the genome of abaca var. Abuab was assembled on scaffold level and annotated, providing a valuable resource for gene discovery (Galvez et al. 2021). More recently, another abaca reference genome was assembled on a chromosome-level (Zhou et al. 2024). On a different study, phylogenetic analysis between *M. balbisiana* and *M. textilis* showed that the evolutionary divergence time was estimated at 7.8 million years ago, using genomic DNA sequencing (Martin et al. 2025).

Despite these developments, studies on the molecular underpinnings of heterosis in abaca, to our knowledge, have not been reported. Unlike model organisms such as *Drosophila* and *Arabidopsis*, and staple crops such as rice and maize, the modes of expression inheritance which drives heterosis have not been studied in this economically important crop. Heterobeltiosis, however, has been reported in banana, *Musa* spp. AAA (Batte et al. 2020).

Allele-specific expression (ASE) imbalance or simply allelic imbalance is now increasingly recognized to explain heterosis. It is the condition in which one of the two alleles in a genome has significantly expressed higher levels of transcripts relative to the other allele. A number of investigations have been performed to elucidate asymmetric expression and heterosis in many organisms, but these are mostly assayed on F_1_ hybrids (e.g. Li et al. 2021; Verta and Jones, 2019; Wittkopp et al. 2004). Studies on hybrid vigor utilizing more advanced hybrid derivatives are scarce. Recently, a study reported the heterotic characteristics of a diploid F_2_ potato population capitalizing on multi-omics approaches (Li et al. 2024). On the other hand, an early study on cucumber demonstrated heterotic effects on fruit weight per plot and fruit number per plot, among hybrids and backcrosses with dominance as the major contributing factor of inheritance (Ghaderi, 1979).

As backcrossing aims to recover the genetic background of one of the parents, it reduces the heterozygosity (or increases the homozygosity) of the hybrids. Heterotic effects, therefore, are expected to be modest in backcross generation. Nevertheless, we posit that the presence of residual heterozygous sites in the backcross genome may still allow interactions between the two alleles contributing to the superior phenotypic performance of this relatively advanced line.

In our previous work on similar genotypes (abaca × banana backcrosses), we concluded that there exists asymmetric expression in BC_2_ and BC_3_ (Ereful et al. 2022b). Hybrid vigor, however, was not investigated that time due to lack of replicates among samples resulting to insufficient statistical power. Since there was allelic imbalance, we hypothesized that expression heterosis is in effect in these backcrosses; thus, this paper. Here, we assessed the role of regulatory interactions (cis and/or trans regulatory factors) in explaining modes of expression performance which drives heterosis. We capitalized on RNA-seq to shed hints on the expression heterosis of BC_2_ (Bandala) and assess its association with regulatory differences between the two interspecifically divergent parental *Musa* species.

## Materials and Method

In a previous project, Abuab, an abaca variety with a high fiber quality but is susceptible to ABTV, was crossed to a virus-resistant wild banana variety, Pacol (*M. balbisiana* Colla). The hybrids were backcrossed several times to Abuab to create BC_1_, BC_2_ (locally named as Bandala), and BC_3_, with selection procedure carried out each generation to identify hybrids with superior fiber quality and resistance against ABTV. BC_2_ hybrids were found to have high fiber quality and resistance against the virus pathogen (Labrador et al. 2020; Mati-om et al. 2024; Parac et al. 2020; Parducho et al. 2020).

Here, we assayed Bandala to investigate heterosis and its relationship with regulatory differences. To easily visualize the workflow implemented in this paper, a schematic diagram is illustrated in Supp. Fig. 1. (The terms ‘BC_2_’ and ‘Bandala’ will be used interchangeably in this paper).

### Sample Collection

Leaf samples were collected from non-stressed 3-month-old *Musa* genotypes – Abuab (Ab), Pacol (Pa), and Bandala (Ban) – from the abaca collection at Feeds and Industrial Crop Section (FICS), Institute of Plant Breeding (IPB), University of the Philippines Los Baños (14°09′09.7″ N, 121°15′39.2″ E) on 3 October 2023, 10 am. Twelve leaf tissues (4 biological reps per variety) taken from the proximal section of young leaves, were snap-frozen in the field and stored at -80°C until further processing.

### RNA Extraction and Sequencing

Total RNA was extracted using a modified CTAB-based protocol as previously described (Ereful et al. 2022a). RNA quality was assessed using a Nanodrop spectrophotometer, with samples showing an A260/280 ratio of 1.9–2.2.

Nine RNA samples (3 reps per variety) were sent to Macrogen, South Korea for RNA-seq using Illumina Novaseq 6000, generating 150-bp paired-end read length. RNA libraries were prepared using the Illumina TruSeq RNA Library Prep Kit with Ribo-Zero rRNA reduction. Briefly, library was prepared by random fragmentation of the cDNA samples followed by ligation and PCR-amplification. All three biological sample–replicates of each genotype were assayed for RNA-seq and showed high correlation and good quality control (QC) stats. Ban rep 2 sample, however, did not pass the initial QC check with succeeding PCA analysis showing its failure to cluster closely with the other two replicates (Supp. Fig. 2). Correlation analysis using Spearman between the other replicates showed relatively low coefficients of 0.87 (between Ban rep 2 and 1) and 0.85 (between Ban rep 2 and 3); rep 1 and 3 showed 0.95 (Supp. Fig. 3). Therefore, Bandala rep 2 was dropped from succeeding analysis, with two reps still retained for Bandala samples.

PCA was performed by transforming the count object into a DESeq object using the variance-stabilizing transformation (‘vst’) function. The relationship of the sample–replicates was eventually visualized using ‘plotPCA’ command in R. Correlation analysis implementing Spearman was performed using ‘cor’ command in R, with raw read counts as the input. Correlation matrix was visualized using the ‘corrplot’ package.

### Bioinformatics pipeline

Quality control of reads was conducted using FastQC (Andrews, 2010). Abuab reads were aligned against the *M. textilis* Abuab reference genome (Galvez et al. 2021); Pacol reads against the *M. balbisiana* (Mba) double haploid – Pisang Klutuk Wulung (DH–PKW) using STAR (Dobin et al. 2013) with number of matched bases set to at least 5 (--outFilterMatchNmin 5), and with allowed maximum number of multiple alignments set at 100 (--outFilterMultimapNmax 100). Both reference genomes were pre-indexed prior to mapping, also using STAR. As Bandala hosts both genomes, we concatenated the Abuab and DH–PKW reference sequences. Reads derived from the Bandala samples were aligned competitively against this merged reference assemblies, using the same tool (STAR) and arguments. BAM files were processed using SAMtools (Li et al. 2009) for downstream analyses.

### Read Counting and Alignment Analysis

Mapped reads were counted using featureCounts (Liao et al. 2013) of the subread package at the gene level (option: gene_id) ensuring only the reads uniquely mapping were counted (option: --primary). In the BC_2_ genome, this represents allele-specific expression after mapping reads competitively in a concatenated reference sequence (Lovell et al. 2018; Kerwin et al. 2020). We added the arguments ‘exon’ as feature type in a GTF annotation to count reads against, applying a minimum mapping quality score of 10 (option: -Q 10) during read counting. Using protein sequences as inputs, orthologs between the two species were identified using OrthoFinder (Emms and Kelly, 2019), retaining only sequences with one-to-one (1:1) correspondence. After ortholog searching, genes from Abuab which matched with multiple DH–PKW genes (one-to-many or 1:m), and vice-versa, were dropped. Their specific allele assignment and asymmetric expression ratios in the BC_2_ hybrid cannot be ascertained.

### Heterosis Analysis

A raw data count of all expressed transcript orthologs across all genotypes (Abuab – BC2 – Pacol) was created. To discern total expression performance of each gene in the backcross hybrid, the Abuab- and Pacol-specific read counts of each transcript ortholog were summed. Orthologs are homologs in different species catalyzing the same reaction (Jensen, 2001). The data count was normalized using median normalization of the DESeq2 package (Love et al. 2014). Pairwise t-test and log (base 2) fold change between two genotypes were calculated to assess statistical and biological significant differences between expression. Pairwise t-test was performed in MS Excel^®^ with 2-tailed distribution and 2-sample unequal variance, or heteroscedastic, since we assume variances are different across sample–replicates. To calculate the Log_2_FC, we computed the average expression of each gene per genotype. In a few cases, we added 1 to avoid 0 denominator. Average normalized read counts with ≥20 combined parental Abuab and Pacol read counts must be satisfied to be considered for further analysis. Modes of expression inheritance were determined as previously reported (Bao et al. 2019; Lovell et al. 2016; Paschold et al. 2012), which can be classified into three “broad classes”: (i) Additive; (ii) Dominance, and (iii) Transgressive. In the subsequent statistical analyses, dominance is more appropriately categorized into two distinct broad Het groups: Lp and Hp dominance.

These were further partitioned into more “specific classes”: Additive, into better-than Abuab and better-than Pacol; Transgressive, into over and underdominance; Dominance, into Low-parent (Lp) and High-parent (Hp) dominance. Abuab serves as the maternal donor in the F_0_ but is the paternal donor in the succeeding backcrosses, thus, it cannot be definitively assigned as maternal or paternal allele; Pacol serves as the paternal donor at F_0_. Therefore, Hp and Lp dominance were further classified as: Hp Ab dom, Lp Ab dom, Hp Pa dom, and Lp Pa dominance. We calculated the log(2) expression ratios between genotypes and plotted them on a 3D plot using ‘plotly’ library in R. Most statistical analyses in this paper were performed in R (v 4.4.1; R Core Team, 2024).

### Regulatory Difference Analysis

We analyzed regulatory differences between the parents and the BC_2_. A raw data frame was created which summarizes the counts of the parental genotypes, Abuab and Pacol, and the Abuab- and Pacol-specific alleles in the Bandala variety. The raw read counts of the parents and the backcross hybrids were normalized using median normalization in DESeq2 (Anders and Hubers, 2010). Only genes which have a total normalized parental read counts of at least 20 were further analyzed for regulatory differences (Ab + Pa ≥ 20) as previously employed (McManus et al. 2010; Shi et al 2012).

Differential expression (DE) between the parents (Ab vs Pa) was performed using DESeq2 (Love et al. 2014) which uses negative binomial modelling of the counts, followed by Wald’s test at FDR-corrected p-value < 0.05 (Benjamini and Hochberg, 1995). Minus–Average (MA) plots were generated using the ‘plotMA’ command with the results (‘res’) of DE as input. DE to asses cis/trans regulatory divergence was traditionally performed using binomial (McManus et al. 2010; Wittkopp et al. 2008). However, due to development in computational biology, we preferred negative binomial in DESeq2 to account for overdispersion in modelling the counts since the variance in RNA-seq datasets is greater than the mean. Likewise, DE between the two specific alleles in the backcross hybrids (Ab- and Pa-specific alleles) was tested using DESeq2 followed by FDR analysis. Transcript orthologs with an FDR < 0.05 is said to be significantly different. Any significant expression difference between the parents and between the two specific alleles in the hybrids is attributed to cis. In a Cartesian plane (Fig. 3), points which represent genes, will lie on the x = y axis.

Any expression performance that could not be explained by cis was ascribed to trans (T = P – H, Fisher, FDR < 0.05) and points will lie on the y = 0. We further classified genes into several categories: (i) if cis–trans regulatory difference resulted to zero expression difference, compensating is in effect and points will lie closely to x = 0 (significant DE in the BC_2_ but not between the parents; significant T); (ii) if cis–trans regulation favor expression of the same allele, synergistic interaction is operating (significant DE in both parents and BC_2_; significant T); (iii) if cis–trans regulation favor the expression of opposite alleles, antagonistic interaction is in force and points will lie closely to y = –x (significant DE in both parents and BC_2_; significant T).

These cis/trans regulatory differences have been adapted from previous papers (McManus et al. 2010; Wittkopp et al 2008; Shi et al 2012) and are described statistically in our github repository site https://github.com/nelcaster7/abaca_heterosis_2025. Ambiguous and conserved are likewise described in our github page.

### Contingency Pearson Chi-squared (χ^2^) test

We performed contingency Pearson χ^2^ test using two data inputs for heterosis (Het): (i) the broad classes, and (2) specific classes (explained above) followed by correspondence analysis using ‘ca’ library in R. This aims to determine association of Counts as response variable to Reg (cis and/or trans regulatory factors or interactions) and Het (additive, dominance, transgressive) as predictors.

Pearson residual analysis was conducted to determine the discrepancies between observed and expected counts and to estimate the contribution of each Reg – Het cell to the χ^2^ statistic. Significance level for each Reg – Het cell was calculated using ‘fisher.test’ function (Fisher exact test; R) in a 2 × 2 contingency table. Cramér’s V (Cramér, 1946) was further estimated to assess the level of association between Reg and Het.

GO enrichment analysis was run using OmicsBox (v 2.1.2; Biobam, 2019) with Cloud BLAST and Cloud InterproScan both included in the workflow. Database search was performed using BLASTx-fast program (Altschul et al. 1990), with non-redundant protein sequences (nr v5) as query, and Musa DB as Taxonomy filter to increase specificity of functional annotation. An e-value of 10^-5^ was implemented.

### Generalized Linear Model (GLM)

A GLM in Poisson and Negative Binomial (NB) regressions was used to model the Counts with the categorical variables, Reg and Het. However, results using GLM could not be concluded as more observations are required to come up with a more robust statistical analysis.

## Results and Discussion

In this paper, we aim to assess expression heterosis in the BC_2_ genome and its association with regulatory differences. In contrast with our previous studies which utilized the central whorl of the pseudostems (Ereful et al. 2022a; Ereful et al. 2022b), here, leaf samples were assayed for RNA-seq to interrogate asymmetrically expressed and heterotic genes associated with photosynthesis and fiber synthesis.

### Reads and Mapping Statistics

FastQC detected no low-quality bases and adapters on the reads, therefore, no pre-processing steps were necessary. Results of the sequencing assay are summarized in Table 1. PCA confirms the close association of genotypic replicates (except Bandala rep 2; Supp Fig. 2). Correlation analysis implementing Spearman showed very strong degree of relationship between genotypic replicates (except for Ban rep 2), with coefficients of at least 0.92 (Supp. Fig. 3).

**Fig. 1.**
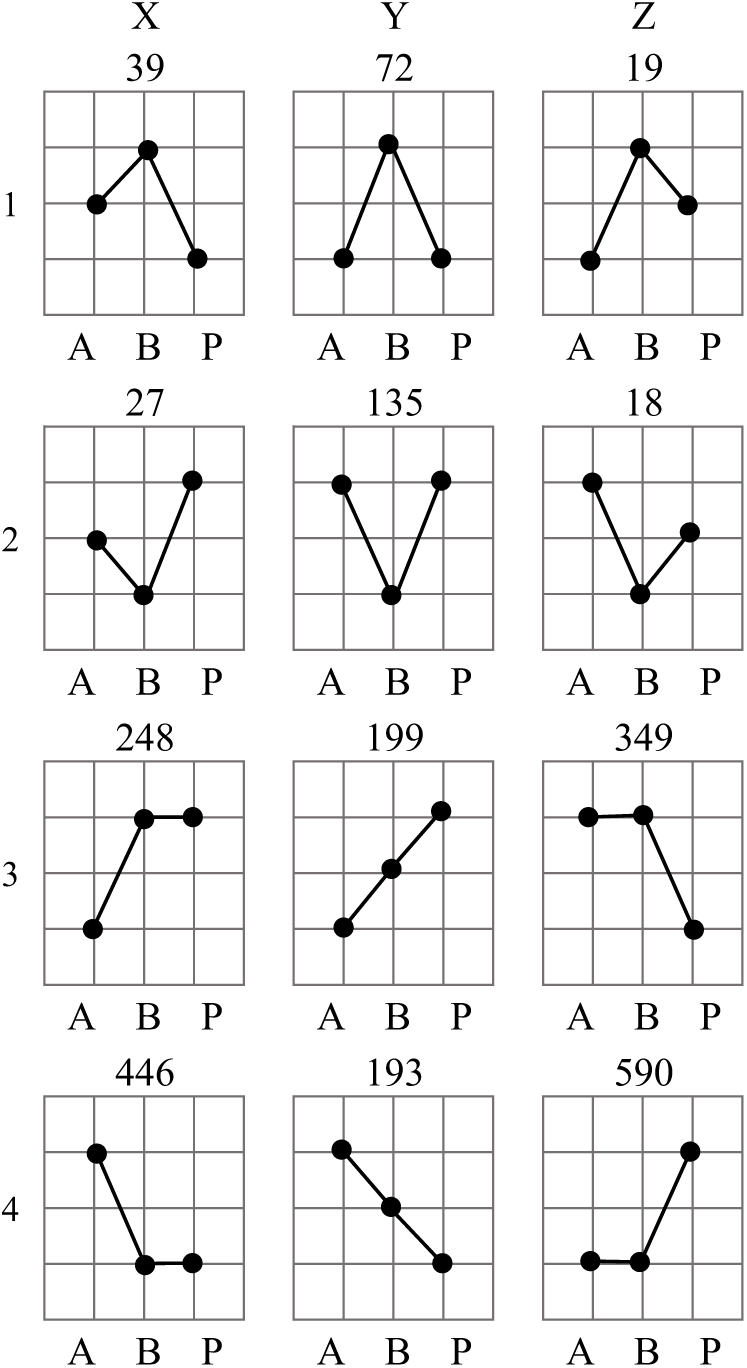
Heterosis in the advance backcross hybrid (BC_2_), Bandala. Two-dimensional grid node demonstrating expression performance of the BC_2_F_1_ Bandala (B) relative to its parents, Abuab (A) and Pacol (P). Values above each panel indicate number of genes (or transcript orthologs) demonstrating the specific type of gene action. Row 1 shows isoforms which exhibits overdominance; 2, underdominance; 3X, Hp Pacol dominance; 3Z, Hp Abuab dominance; 4X, Lp Pacol dominance; 4Z, Lp Abuab dominance; 3Y (better-than Abuab) and 4Y (better-than Pacol), mid-parent expression performance or additive.

**Fig. 2.**
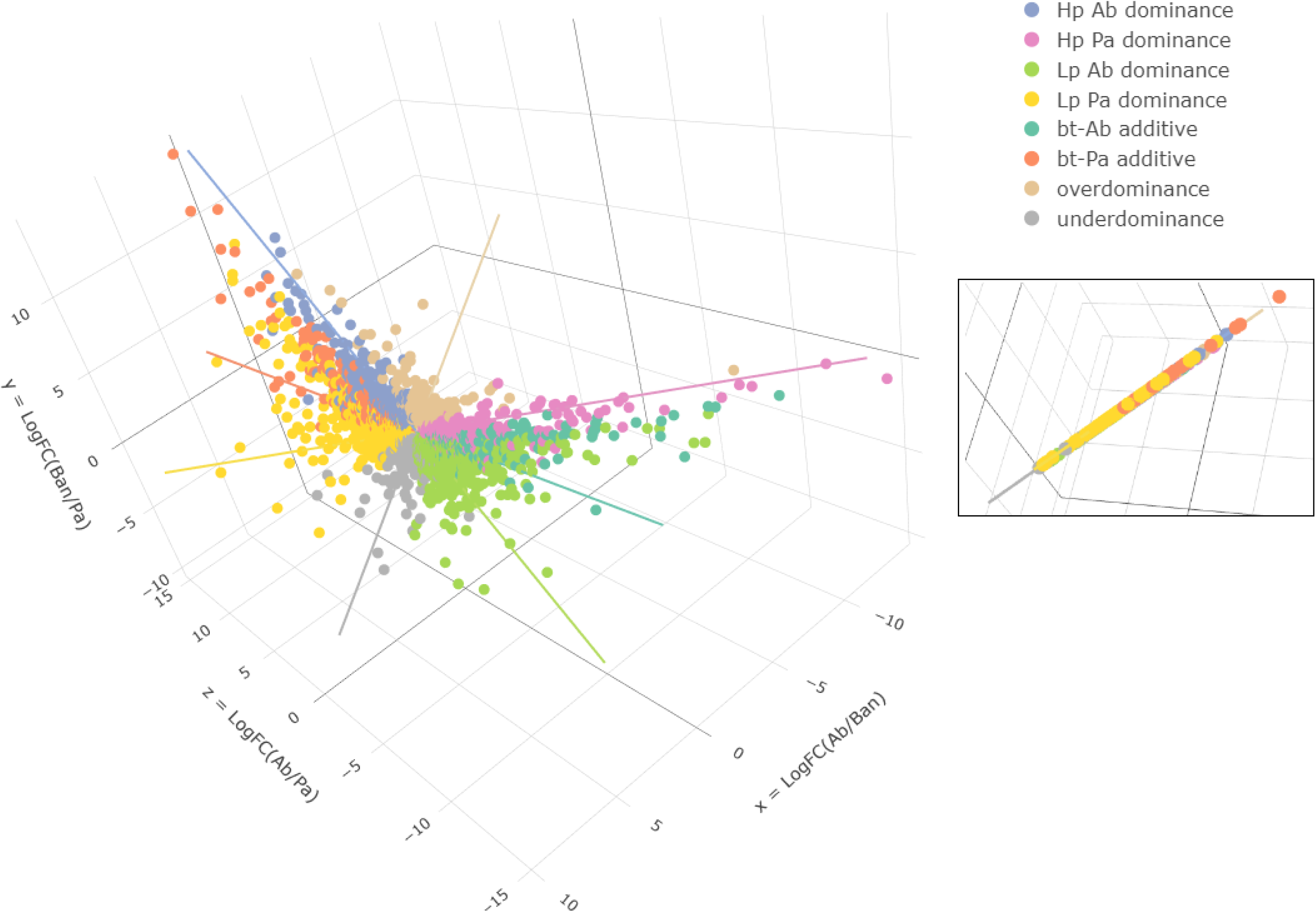
The heterosis architecture of BC_2_ (Bandala), depicted in a 3D space. Gene–points belonging to a specific mode of inheritance coincide closely to a specific line which is described by an equation (Table 3). Inset: at a particular angle, heterotic gene–points lie in a tilted flat line (see our github page to view the 3D movie file: https://github.com/nelcaster7/abaca_heterosis_2025). (Legend: Hp, High-Parent; Lp, Low-parent; Ab, Abuab; Pa, Pacol; bt, better-than).

**Figure 3.**
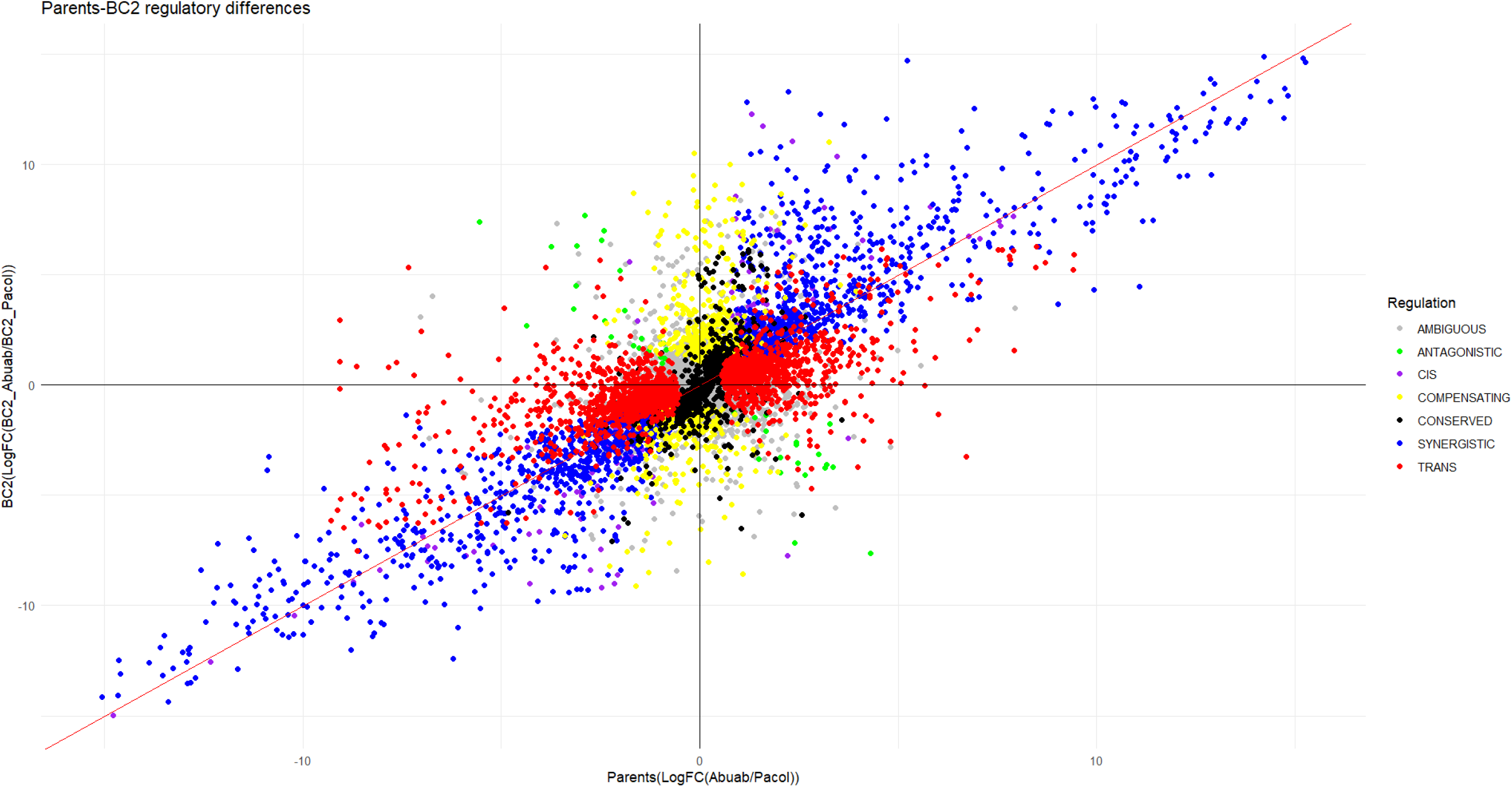
The cis/trans regulatory difference landscape between the parents (Ab/Pa) and BC_2_ abaca, Bandala.

**Table 1.** Number of reads and percentage mapped of the parents and their BC2 hybrid.

| Sample replicate | Number of PE reads (×2) | Q20(%) | Q30(%) | % Mapped |
| --- | --- | --- | --- | --- |
| Abuab rep 1 | 63951106 | 97.7 | 93.7 | 94.13 |
| Abuab rep 2 | 59565184 | 97.8 | 94.0 | 90.10 |
| Abuab rep 3 | 76042328 | 97.8 | 94.0 | 95.37 |
| Bandala rep 1 | 75713368 | 97.7 | 93.9 | 97.40 |
| Bandala rep 2 | 75555304 | 98.4 | 95.3 | 94.94 |
| Bandala rep 3 | 65734442 | 97.9 | 94.2 | 97.79 |
| Pacol rep 1 | 66126688 | 97.8 | 94.1 | 98.31 |
| Pacol rep 2 | 73111314 | 97.8 | 94.1 | 98.93 |
| Pacol rep 3 | 74513968 | 97.8 | 94.1 | 99.10 |

Quality analysis of the reads showed Q20 and Q30 of at least 97.7% and 93.7%, respectively, suggesting high quality reads were obtained. Mapping percentage showed at least 90% indicating high alignment rates. MA plots between the parents (Supp. Fig. 4) and between the parent-specific alleles in BC_2_ (Supp. Fig. 5) showed symmetrical data cloud indicating that normalization was effective with respect to library size.

## Heterosis

We calculated the pairwise log-transformed ratios of the normalized expression counts of each gene, then classified the ratios based on different modes of expression inheritance (see Materials and Method; results are summarized in Table 2 and illustrated in Fig. 1). 10,939 genes with 1:1 correspondence were detected between *M. textilis* (abaca var Abuab) and *M. balbisiana* (var DH – PKW) using OrthoFinder. This is nearly one-third of the abaca genome. Hence, two-thirds of the genes have either 1:m (vice-versa) correspondence or do not have counterpart (orphan) gene and cannot be assigned to a particular ortholog.

**Table 2.** Number of genes classified in each broad and specific heterosis classes based on expression performance. (Letters A, B, and P in column 3 stand for Abuab, Bandala or BC2, and Pacol, respectively).

| Broad classes | Specific heterosis classes | Notation | Number of genes |
| --- | --- | --- | --- |
| Additive | Better-than Pacol | $A > B > P$ | 193 |
| | Better-than Abuab | $P > B > A$ | 199 |
|  | Total Number of Additive genes |  | 392 |
| Dominance | Lp Ab dominance | $P > A=B$ | 590 |
| | Lp Pa dominance | $A > P = B$ | 446 |
| | Hp Ab dominance | $A = B > P$ | 349 |
| | Hp Pa dominance | $P = B > A$ | 248 |
|  | Total number of Dominant genes |  | 1633 |
| Transgressive | Overdominance (>Hp) | $B > A = P; B > A > P;$<br>$B > P > A$ | 130 |
| | Underdominance (<Lp) | $A = P > B; A > P > B;$<br>$P > A > B$ | 180 |
|  | Total number of Transgressive genes |  | 310 |

**Table 3.** General and specific line equations corresponding to broad and specific Het classes, respectively, which describe genes clustering in each mode of inheritance.

| Broad Het classes | Specific Het classes |
| --- | --- |
| Transgressive: $x + y = 0$ ($z = 0$) | if $x < 0$: Overdominance |
| | if $x > 0$: Underdominance |
| $P_1$ (Hp and Lp) dominance: $y - z = 0$ ($x = 0$) | if $y > 0$: $P_1$ Hp dominance |
| | if $y < 0$: $P_1$ Lp dominance |
| $P_2$ (Hp and Lp) dominance: $x - z = 0$ ($y = 0$) | if $z < 0$: $P_2$ Hp dominance |
| | if $z > 0$: $P_2$ Lp dominance |
| Additive or mid-parent: $x - y = 0$, $2x - z = 0$ | If $z < 0$ better-than $P_1$ |
| | if $z > 0$ better- than $P_2$ |

There were 9457 genes found to have a combined normalized parental read counts of more than 20 (Ab + Pa ≥ 20). Of these, 4662 showed no DE which means that the comparative level of expression of genes among the genotypes are not significantly different (Ab = Ban = Pa; pairwise t-test, P ≥ 0.10). These genes are relatively equally expressed across the genotypes and presumably do not contribute to hybrid vigor; hence, non-heterotic.

Further analysis showed that 2,459 transcript orthologs cannot be assigned to a particular heterosis class due to statistical conflicting patterns (i.e., ambiguous). This leaves only 2336 genes which can be unambiguously assigned to a specific mode of expression inheritance (pairwise t-test, P<0.10). The use of a lenient pval < 0.10 has been implemented previously (Ereful et al. 2021) to accommodate broader number of transcript orthologs. We classified these genes based on their inheritance mode (Table 2) and are illustrated in Fig. 1 using 2D grid node for easier visualization. (For list of genes and their respective mode of expression inheritance, see Supp. Table 1).

If we are to classify the genes based on the broad Het classes, dominance emerged as the preferred mode of expression inheritance (1,633 genes). This significantly surpassed the number of additive (392) and transgressive (310) genes. Apparently, 83.2% of these heterotic genes were non-additive. This finding is similar to Li et al.’s report in which 86% of the hybrid gene expressions in cabbage were non-additive (Li et al. 2021).

Dominance, if classed based on the level of expression, showed 1,036 and 597 transcript orthologs for Low-parent (Lp) and High-parent (Hp) dominance, respectively. Notably, the combined parental Lp dominant genes exhibited significantly higher prevalence compared to Hp dominant genes (binomial exact test of equal proportion, P < 0.001).

If partitioned based on allelic preference, Ab dominance exhibited a significant preference over Pa dominance, with 924 and 702 genes, respectively (binomial, P = 4.0e-08, at p = 0.5). However, this could be ascribed to the larger genetic presence of Ab over Pa. If we are to use p = 0.875 (due to 87.5% Ab and 12.5 Pa alleles in BC_2_), then there is significant deviation of the observed gene counts over the expected counts (binomial, P < 2.2e-16).

On the other hand, we identified 130 genes showing overdominance and 180 genes exhibiting underdominance. Here, underdominant significantly supersedes overdominant genes (binomial, P = 0.005). The role of underdominance in hybrids has been under recent scrutiny. It was previously identified as the predominant heterosis class associated with growth advantage in *Arabidopsis thaliana* F_1_ hybrids (Yuan et al. 2023).

### Heterotic and non-heterotic genes

A large number of genes were non-heterotic (4,662) demonstrating their equally significant expression across the three genotypes. For example, the gene LOCUS_008714 encoding for Phenylalanine ammonia-lyase (PAL) is non-heterotic. It catalyzes the initial reaction in phenylpropanoid pathway, providing precursors for all downstream pathway including lignin biosynthesis (Xie et al. 2018). Additionally, the photosynthesis genes LOCUS_017803 and LOCUS_022009 which encode for Photosystem I chlorophyll a/b-binding protein and Photosystem I reaction center subunit V, respectively, were both non-heterotic.

In our previous work, the fiber synthesis-associated gene (LOCUS_024375) which encodes for cellulose synthase (putative) was found up-regulated in abaca pseudostem (Ereful et al. 2022a). In the current study, the gene was identified to be dominant (i.e. Hp Ab dom) making it an interesting feature for future study. Additionally, we detected eight non-heterotic and four Hp Ab dominant genes encoding for either cellulose synthase or its catalytic subunit (Supp. Table 1). Both lignin and cellulose are major components of abaca fiber.

### Modes of expression inheritance in 3D plot

The pairwise log (2)-transformed ratios of the normalized expression counts of each gene were plotted in a 3D plot (Fig. 2). The dots, representing heterotic genes, lie on a tilted 2D straight plane in a 3D space due to the transitive relationship of the log-transformed ratios (Fig. 2 inset). That is, the ratios have been computed in a pairwise fashion or cyclic manner among the three genotypes, i.e., Ab/Ban, Ban/Pa, and Ab/Pa, where the pairwise log (base 2) of each expression ratio is assigned as x, y, and z, respectively. Each gene–point on the plot follows the equation x + y = z or log_2_(P_1_/H) + log_2_(H/P_2_) = log_2_(P_1_/P_2_), where P_1_ = parent 1; P_2_ = parent 2; and H = hybrid. In our case, this is equivalent to log_2_(Ab/Ban) + log_2_(Ban/Pa) = log_2_(Ab/Pa).

Interestingly, the plot showed that genes belonging to a specific mode of expression inheritance tend to cluster closely in space. For example, overdominant genes are directed from the origin (0,0,0) towards (–x, y, 0) region with an implicit form of the line equation: x + y = 0 (z = 0; x < 0) (indicated by the brown line in Fig. 2). This region hosts genes that are significantly expressed in the hybrid as compared to the parents. The interaction between cis and trans resulted to the expression amplification of the focal gene (explained further in the next section). On the other hand, underdominant genes are concentrated on the (x, –y, 0) region of the 3D space (gray line in Fig. 2) and has the same equation as overdominance, where x > 0. Allelic interactions in the BC_2_ resulted to the down-regulation of the orthologs compared to the parents.

On the other hand, genes classified as Hp Ab dominant occupy the (0, y, z) zone with the equation form: y = z (x = 0, y > 0) (blue line in Fig. 2). In terms of expression, this line (Hp Ab dominance region) coincides with genes in Bandala which are expressed in the same level as that of genes in the Abuab but are lowly expressed in Pacol (i.e. Pa < Ab = Ban). Lp Ab dominance has the same line equation except that y < 0. As Abuab possesses high fiber quality traits (not Pacol), future breeding efforts are directed towards selecting features exhibiting Hp Ab dominance and overdominance associated with high fiber quality; thus, this interest in overdominant and Hp Ab dominant genes. Early and recent papers have hypothesized these two gene actions to drive heterosis (reviewed in Paril et al. 2024; Yu et al., 2021).

The finding that genes belonging to the same mode of inheritance aggregating closely in a 3D plot is consistent with our previous work on heterosis in rice F_1_ hybrids exposed to non- and water-stress conditions (Ereful et al. 2021; see Supp. Fig. 6 and 7 for non- and water-stress conditions, respectively; supplementary figures were unpublished). Similar to the current paper, the plots contain lines with the same equations as Bandala’s, with gene–points lying on the same 3D plot: x + y = z, which corresponds to log_2_(IR64/Hyb) + log_2_(Hyb/Apo) = log_2_(IR64/Apo), where IR64 and Apo are parental indica rice varieties; Hyb, hybrid. The fitted line equations of each broad and specific Het classes of expression inheritance are, likewise, the same as Bandala’s (summarized in Table 3).

These line equations were consistent between the two different taxa (*Musa* vs rice) assayed along with their respective progenies (abaca BC_2_ and rice F_1_). These were, likewise, consistent between the two intra-sub-specifically related rice varieties (i.e. IR64 and Apo) exposed under two contrasting environmental regimens (non- and water-stress conditions).

These line equations may exhibit universality awaiting further validation through further research across other crops. It may have pending applications in predicting modes of expression performance, thereby aiding future selection strategies.

### Phenotypic heterosis

In our separate but concurrent work on abaca, we observed overdominance of the BC_2_ over its parents on two economically important morpho-agronomic traits – fiber length and tensile strength. BC_2_ exhibited 275 cm fiber length while its parents, Ab and Pa, measured 250 and 150 cm, respectively. On tensile strength, the hybrid Bandala, and its parents Ab and Pa exhibited 40.3, 38.1, and 31.2 kg·force/g/m (all phenotypic values are average of 12 suckers per genotype with data collected from 2018 to 2022; analyzed using LSD). Here, the BC_2_ is outperforming its parents.

On the other hand, Ab, Pa, and Bandala showed plant heights (cm) of 273, 321, and 290, respectively, to which the hybrid is exhibiting midparent phenotypic performance. Stem Fresh Wt. exhibited Hp Pacol dominance with Abuab, Pacol, and Bandala weighing 25.0, 29.0, and 30.3 kg, respectively.

Apparently, the backcross hybrid exhibits various modes of inheritance both on the transcriptomic and phenotypic levels.

## Regulatory differences

Analysis of regulatory differences was carried out to assess variations in the cis and/or trans regulation between the parents and the hybrids. In several investigations, regulatory difference was estimated between parental genotypes and their F_1_ hybrids (e.g. Wittkopp et all 2004; Wittkopp 2008; McManus et al. 2010). In this study, materials include backcrosses, therefore, the accumulated regulatory differences have been shaped by evolutionary divergence between *M. textilis* and *M. balbisiana* (Martin et al. 2025; Zhou et al. 2024) and repeated backcrossing and selection process.

To depict the cis and/or trans regulatory architecture, Fig. 3 was created to demonstrate the suite of cis/trans regulation type. Results of this analysis are summarized in Table 4 showing the different regulatory factor/interaction classes and the corresponding number of genes (see Supp. Table 2 for cis and trans regulatory assignment of each gene; see Materials and Method for complete statistical analysis; adapted from McManus et al. 2010; Shi et al 2012; Wittkopp et al 2008).

**Table 4.** Number of genes exhibiting each cis and/or trans regulatory divergence.

| Regulatory factor / interaction | Number of genes |
| --- | --- |
| cis | 162 |
| cis + trans (synergistic) | 1696 |
| cis × trans (antagonistic) | 41 |
| compensating | 703 |
| trans | 1839 |
| ambiguous | 3875 |
| conserved | 993 |

Overall, the combined effect of trans and cis + trans (synergistic) interactions largely explained and contributed to regulatory differences and successive selection biases, showing 1,839 and 1,696 genes, respectively, with a combined percentage of ∼80% of the total number of genes (ambiguous and conserved were dropped from the analysis). Goodness-of-fit Chi-squared test (χ^2^ = 3,193, *P*<10^-5^) of the data strongly significantly indicates a non-neutral regulatory difference (against the H_0_ of equal contributions). Synergistic interaction and trans are largely in effect, explaining the selection bias towards the beneficial traits. Suckers with superior fiber quality were preferentially selected demonstrating a directional selection. This happens when one of the two extreme ends of the population is favored. These results suggest that directional selection leads to enhancing cis and trans effects (Pearson residual for synergistic = 27.1; 23% contribution to the χ^2^ statistic). Note that enhancing (Bao et al. 2019; Shi et al. 2012) and synergistic (McManus et al. 2010; Wittkopp et al. 2008) are synonymous in which cis and trans act in the same direction in regulating gene expression.

On the other hand, only 41 genes exhibited antagonistic interaction (cis × trans). These findings suggest that the combined divergence and successive selection procedures favor the accumulation of synergistic alleles while suppressing antagonistic alleles. The modest number aims to potentially mitigate the undesirable effects of opposing gene interactions (Pearson residual = -28.43; 25.31% contribution). Antagonism has been reported to contribute to misexpression in hybrids, thus, hybrid incompatibility (McManus et al. 2010).

The finding that trans and cis + trans largely explain parents – BC_2_ regulatory differences is contrary to our previous study (Ereful et al. 2022b) in which cis explains mostly the regulatory difference. These discrepancies may be partly ascribed to the different tissues analyzed where we previously assayed the inner whorl of abaca pseudostems. This was also observed in maize and teosinte in which three tissues (ear, leaf, and stem) showed changing proportions of cis and trans genes exhibiting dominant or additive inheritance (Lemmon et al. 2014). Furthermore, leaves are directly exposed to both biotic and abiotic factors including photosynthesis. This may explain the accumulation of trans factors known to participate in response to environmental treatments (Tirosh et al. 2009). On the other hand, the central inner core of the abaca pseudostems are thickly insulated and are largely engaged in fiber synthesis. It must be noted, though, that results derived by comparing these two studies must be interpreted with caution as materials are not collected in the same year and season and therefore are not exposed to the same environmental conditions.

Cis-regulatory factor played a limited role in explaining these differences, which is likely ascribed to the near-homozygosity of the backcross to one of the parents. This aligns with our previous study demonstrating that repeated selection and backcrossing affects cis-regulatory elements by increasing homozygosity (or reducing heterozygosity) with each advancing generation, i.e. from BC_2_ to BC_3_ (Ereful et al. 2022b).

## Regulatory differences drive heterosis

Recent papers have reported the influence of regulatory changes to modes of expression inheritance (Bao et al. 2019; Ereful et al. 2021). In this study, we tested the association between the predictors regulatory difference (Reg) and heterosis (Het) by classifying each gene according to regulation type (cis and/or trans regulatory factor or interaction) and the heterosis class they exhibited. Results are tabulated in Table 5 with Het being categorized into specific classes of mode of inheritance (also described in Materials and Method).

**Table 5.** Number of genes and their regulation types, categorized into specific Het classes. The number of genes for each Regulation type under Transgressive, Dominance (Lp and Hp Dominance), and Additive were summed to determine broad Het classes.*

| Regulation | Transgressive |  | Dominance |  |  |  | Additive |  |
| --- | --- | --- | --- | --- | --- | --- | --- | --- |
|  | Underdom | Overdom | Lp Pa dom | Lp Ab dom | Hp Pa dom | Hp Ab dom | btr-thn Ab | btr-thn Pa |
| cis | 1 | 0 | 2 | 12 | 22 | 29 | 5 | 2 |
| cis + trans | 4 | 23 | 130 | 210 | 127 | 156 | 154 | 153 |
| cis × trans | 1 | 1 | 5 | 4 | 0 | 5 | 0 | 1 |
| compensating | 14 | 26 | 11 | 18 | 5 | 12 | 0 | 1 |
| trans | 15 | 10 | 178 | 238 | 46 | 84 | 37 | 36 |
| ambiguous | 105 | 57 | 104 | 71 | 38 | 56 | 1 | 0 |
| conserved | 36 | 13 | 3 | 21 | 10 | 7 | 2 | 0 |

432 \* Underdom, underdominance; Overdom, overdominance; Lp, Low-parent; Hp, High-parent; Pa, Pacol; Ab, Abuab; btr-thn, better-than

A contingency Pearson χ² test using the data information (Table 5), coupled with correspondence analysis (CA) was performed to assess the association of Reg and Het to the Counts (Counts ∼ Reg + Het) (ambiguous and conserved were not included in this analysis). The use of specific Het classes aims to capture fine-scale and genotype-specific variations within the Het categories. The results revealed that Counts are not independent of Reg and Het (χ² =561.2, df = NA, P = 10^-5^; based on Monte Carlo simulation with 10⁵ replicates) and that the observed differences are not due to random chances. This finding confirms the association between cis and/or trans regulatory differences and heterosis. This is consistent with recent investigations in which regulatory differences is significantly associated with heterosis (Bao et al. 2019; Ereful et al. 2021).

To visualize the association between Reg types and specific Het classes, a CA factor map was generated (Fig. 4A). Apparently, there appears to be a distinct partitioning of Hp and Lp dominance suggesting a specific regulatory type is in effect. Hp dominance is closely associated with cis and cis + trans; Lp dominance, with trans. The contrasting regulatory mechanisms at play and their dissociation in the CA factor map suggest separating Hp and Lp dominance into two separate categories (Fig. 4B), in contrast with previous papers combining them into a single dominance mode category (Bao et al. 2019; Ereful et al. 2021).

**Fig. 4.**
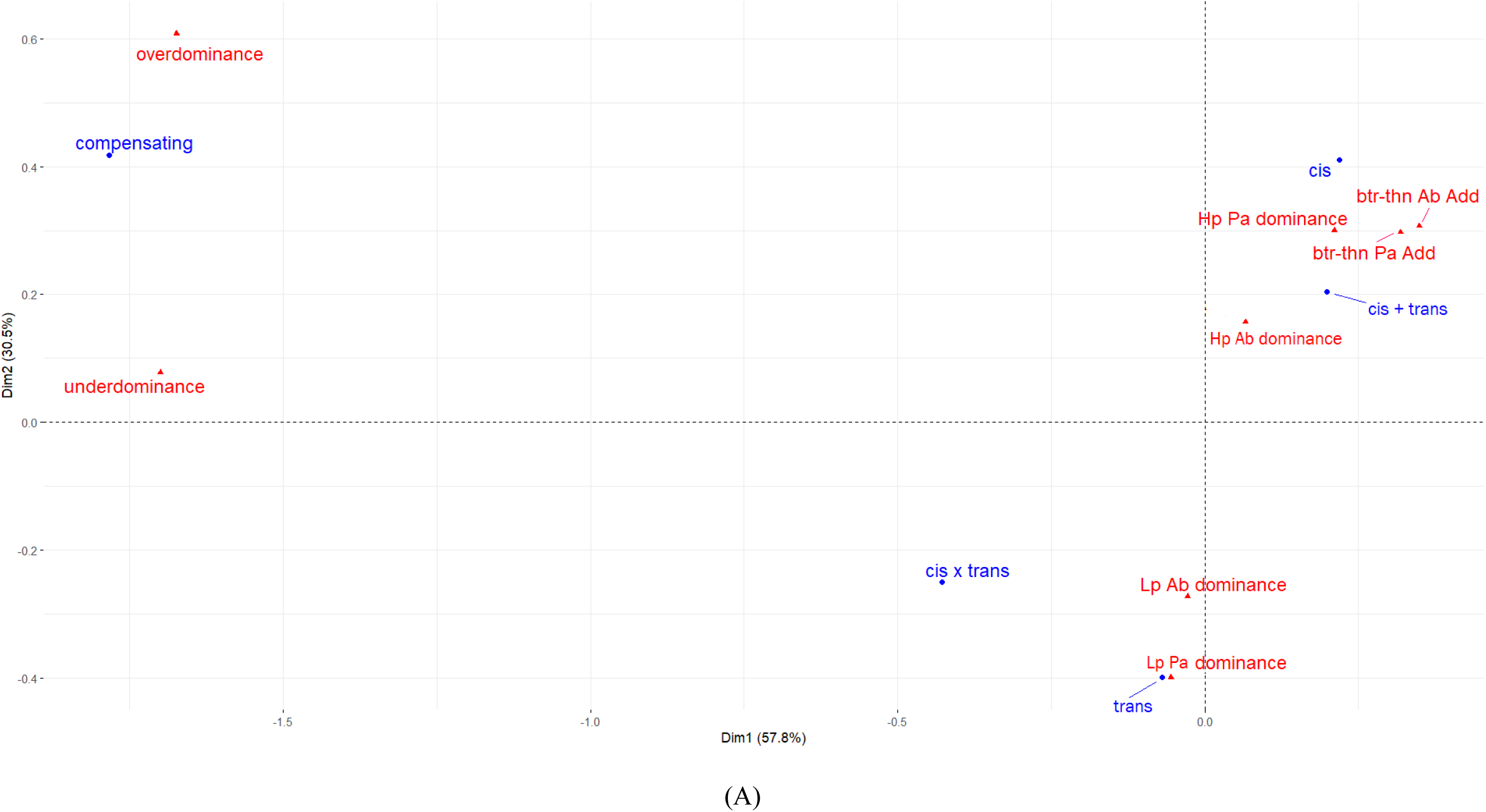
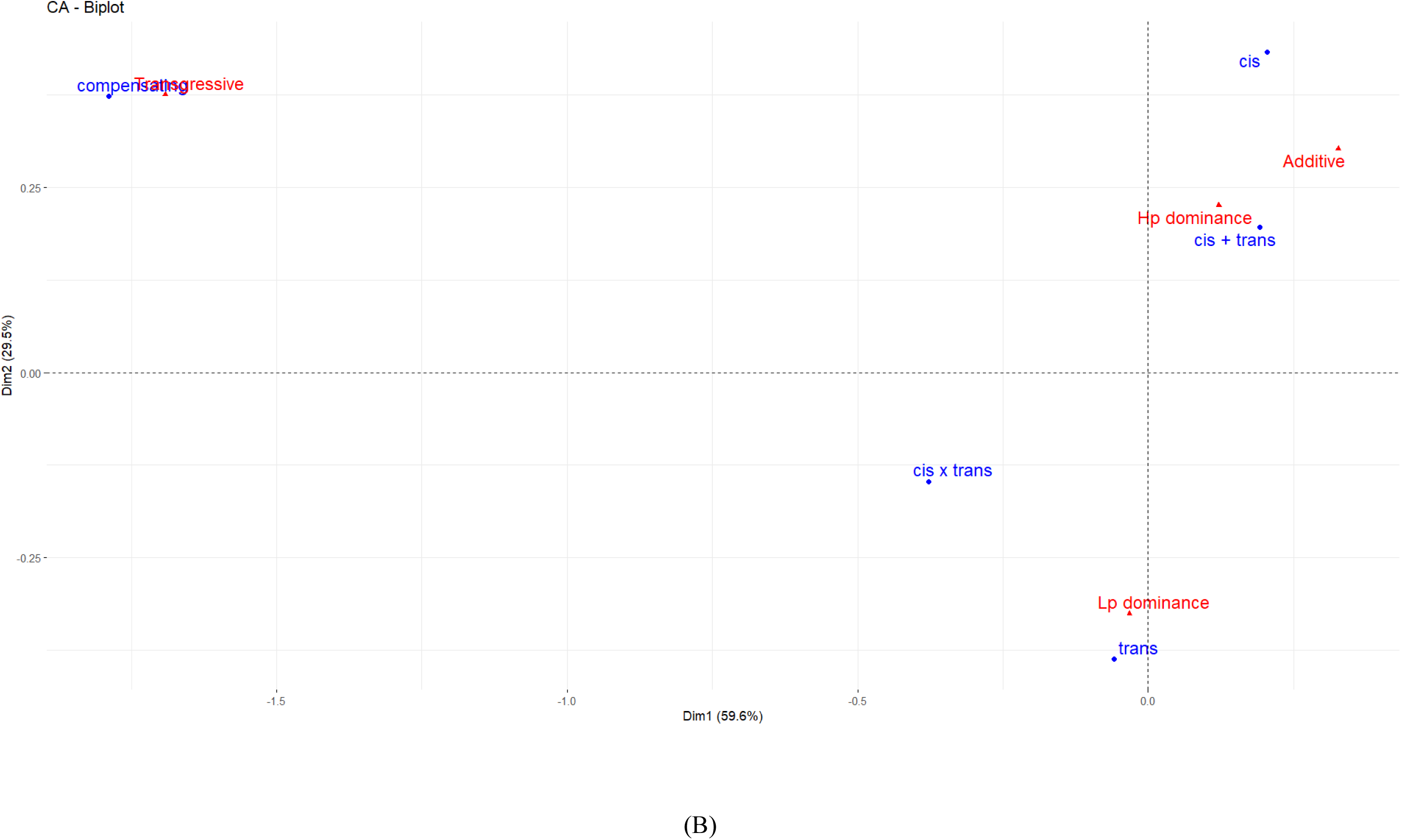
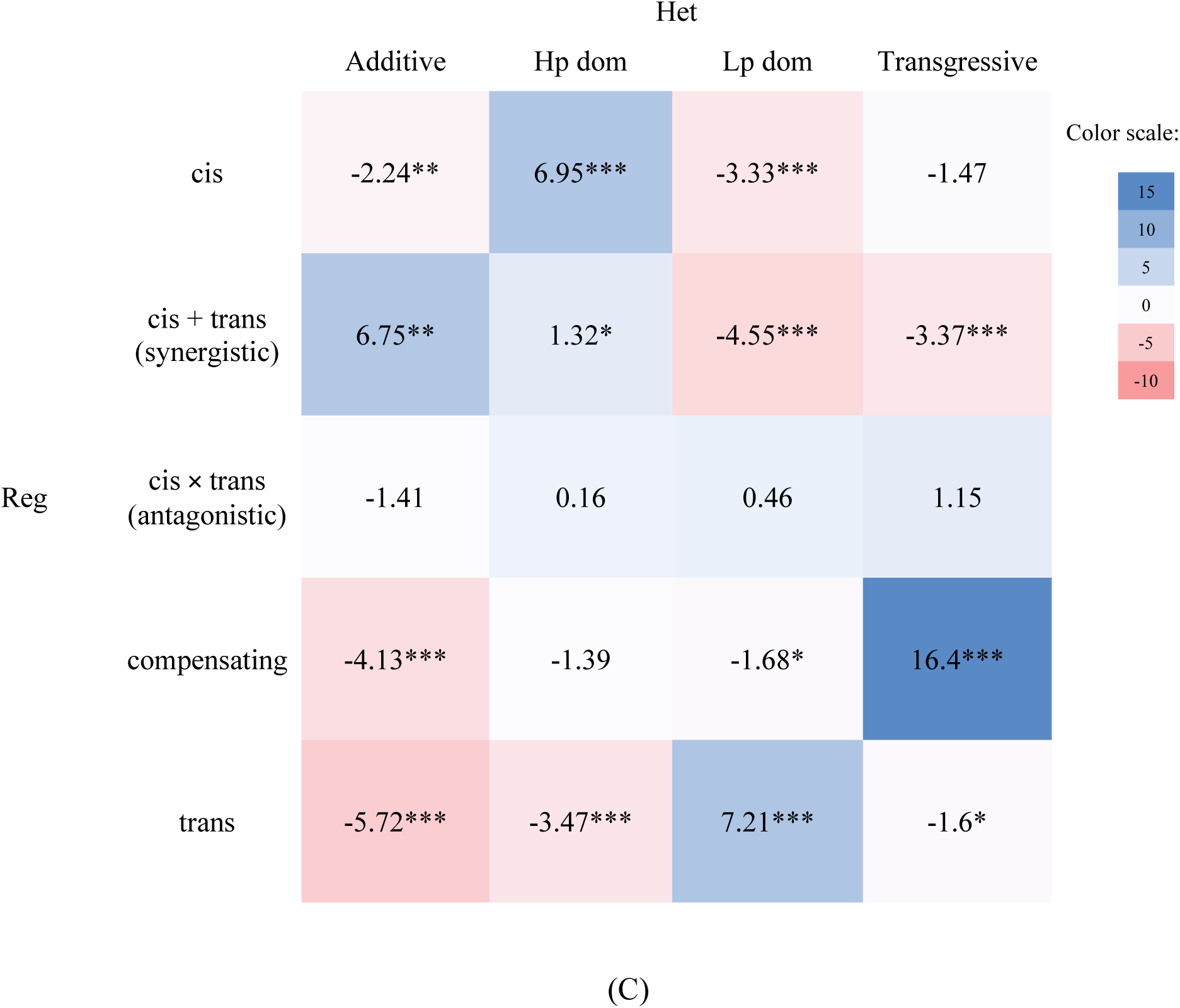
Each Reg type and Het class were plotted on CA factor maps based on (A) specific and (B) broad Het classes. (C) Enrichment–depletion cells were created using contingency Pearson χ^2^ test of independence. Blue cell represents enriched Reg – Het combination; red, if depleted (see Results and Discussion). Numerical values represent Pearson residuals with significance level calculated using Fisher exact test (\**P*<0.05, \*\**P*<0.01, \*\*\**P*<0.001).

The transgressive subclasses, over and underdominance, were both tightly linked with compensating interaction, consistent with recent papers (Bao et al. 2019; Gao et al. 2022). ‘Better-than Abuab’ and ‘better-than Pacol’ are appropriately grouped collectively as Additive since their patterns are closely related and suggest tight association with synergistic (cis + trans) and cis (Fig. 4B and 4C) (further discussed below).

### Pearson residuals

To capture general patterns of heterotic effects, we now partitioned the gene actions into four broad Het classes. CA factor map indicates that the four modes are clearly separated with x-axis (Dim 1) explaining 59.6% of the variance; Dim 2, 29.5% (Fig. 4B). Integrating CA and Pearson residual analysis (Fig. 4B and 4C, respectively) showed that Hp dominance is significantly associated with cis and moderately with cis + trans; Lp dominance, with trans; transgressive, with compensating; and additive, with cis + trans.

In Pearson residual analysis, the contribution of each cell (or the Reg factor – Het class combination) to the χ^2^ statistic is assessed. We say that a Het class is significantly enriched (or overrepresented) if the number of observed gene counts exhibiting a particular Reg type is higher than the expected count; depleted (underrepresented), if the observed count is lower than the expected.

Results suggest that there is significant enrichment of genes exhibiting compensatory interaction under the transgressive mode, with the highest percentage contribution of 49.8% to the χ^2^. This agrees with previous papers in which compensating regulation is enriched for transgressive inheritance in cotton (Bao et al. 2019) and lotus (*Nelumbo nucifera*) (Gao et al. 2022). Compensating alleles exhibit opposing interactions between cis and trans resulting to no expression difference between species in F_1_ (McManus et al. 2010). This balancing effect may have been lost in the backcrosses potentially due to the unequal proportion of the parent-specific alleles. This discrepancy may have resulted to over- or under-expression of alleles resulting to transgressive action, the largest contributor to hybrid vigor in our case. GO enrichment analysis of compensatory genes overrepresented under transgressive mode showed significant association with binding (ion, nucleotide, protein, etc.; Molecular Function (MF); Supp. Fig. 8), and metabolic and biosynthetic processes (Biological Process (BP); Supp. Fig. 9).

Further analysis using Pearson residual revealed that genes diverging in cis are enriched under Hp dom but are depleted in Lp dom. On the contrary, trans genes are enriched in Lp dom but are depleted in Hp dom (Fig. 4C). These findings further underscore the contrasting regulatory mechanisms in effect between these two dominance categories. Cis genes enriched under Hp dom contributed 8.95% to the χ^2^; trans under Lp dom, 9.64%.

Synergistic (cis + trans) genes are significantly enriched under the additive mode and contributes 8.44% to the χ^2^. The interaction between cis and trans resulted to intermediate expression of genes reflecting additivity. No significant enrichment nor depletion of antagonistic interaction was detected under any mode of inheritance possibly due to the low number of counts. Consistent with our observation on regulatory difference (discussed above), antagonistic interaction may have been selected against in favor of more beneficial allelic interactions under directional selection.

*Cramér’s V*. Analysis using Cramer’s V gives a value of 0.318 (χ^2^ = 539.47; n = 1778) if we are to partition Reg into five types and Het into four broad classes (5 × 4 contingency table). This value shows strong relationship between Reg and Het. However, epigenetics (Duarte-Aké et al. 2023), environment (Virmani et al. 1982), and other factors can also influence this cline of gene actions.

In our previous paper on rice (Ereful et al. 2021), the effect of the environmental conditions (Env) showed moderate association with heterosis (Cramér’s V ≈ 0.233) while it has a strong association with regulatory divergence (Reg × Env; V ≈ 0.359). Finally, regulatory divergence in rice hybrids has moderate influence on Het (V ≈ 0.288).

## Conclusion

Taken all together, this study demonstrated that cis and/or trans regulatory factors and their interactions with genes influence hybrid performance. The combined evolutionary divergence and backcrossing–selection procedures have effected allelic imbalance in the backcross hybrids conferring the progenies transcriptional expression versatility. This leads to the repertoire of modes of expression inheritance which propels heterosis. We wish to highlight the need for further exploration of heterosis on more advanced filial and backcross generations to understand the molecular mechanisms of allelic interactions, capitalizing on the rapidly changing sequencing landscape.

## Compliance with Ethical Standards

We adhered to the highest local and international ethical standards on the use of plant materials. Abaca and banana samples collected in this paper are not endangered nor at risk of extinction.

## Conflict of Interest

The authors declare no conflict of interest.

## Data Availability Statement

All datasets are available in EMBL-EBI ArrayExpress (www.ebi.ac.uk/arrayexpress) with an assigned accession number: E-MTAB-14954.

## Funding

This project was funded by the UPLB Basic Research Program of the University of the Philippines Los Baños, Laguna with Project ID 18974.

## Acknowledgment

We used AI-driven tool particularly ChatGPT to assist us in generating 3D plot (Fig. 2) and CA factor map (Fig. 4A and 4B). AI-driven outputs were verified and analyzed by the authors. We would like to acknowledge Dr. Deziree Labrador for her assistance in the field collection of abaca samples. We thank Joonas Kalda of Tallinn University of Technology, Estonia for his assistance on the mathematical analysis. We also thank the members of staff of the Biochemistry – Analytical Services Laboratory, Crop Physiology, and Genetics of IPB, UPLB for additional assistance on the molecular biology part.

## Author Contributions

NCE conceptualized the project, sought funding, analyzed the data, and wrote the manuscript; RCSA performed field sample collection and molecular laboratory procedure; ACL generated the backcrosses and managed the abaca field collections.

## Supplementary Information

Supp. Fig. 1. Schematic workflow diagram of the methodology

Supp. Fig. 2. Principal Component Analysis (PCA) of the normalized RNA-seq datasets of the three genotypes

Supp. Fig. 3. Correlation analysis implementing Spearman between sample–replicates

Supp. Fig. 4. MA plot between Abuab and Pacol varieties

Supp. Fig. 5. MA plot between Abuab- and Pacol-specific alleles in BC_2_ Bandala

Supp. Fig. 6. 3D scatter plot of genes exhibiting various modes of expression inheritance in rice F_1_ hybrid under non-stress condition

Supp. Fig. 7. 3D scatter plot of genes exhibiting various modes of expression inheritance in rice F_1_ hybrid under water-stress condition

Supp. Fig. 8. GO distribution of compensatory genes significantly enriched under transgressive mode of inheritance associated with MF. Figure generated and analyzed using OmicsBox

Supp. Fig. 9. GO distribution of compensatory genes significantly enriched under transgressive mode of inheritance associated with BP. Figure generated and analyzed using OmicsBox

Supp. Table 1. Specific and broad heterosis classification of each gene in the BC_2_ Bandala genotype

Supp. Table 2. Regulation classification of each gene in the BC_2_ Bandala genotype

## Notes

### Competing Interest Statement

The authors have declared no competing interest.

