## Supplementary figures and images for "Molecular Heterosis in the Abaca (*Musa textilis* Née) BC_2_ Hybrid, Bandala"

### Supp. Fig. 1

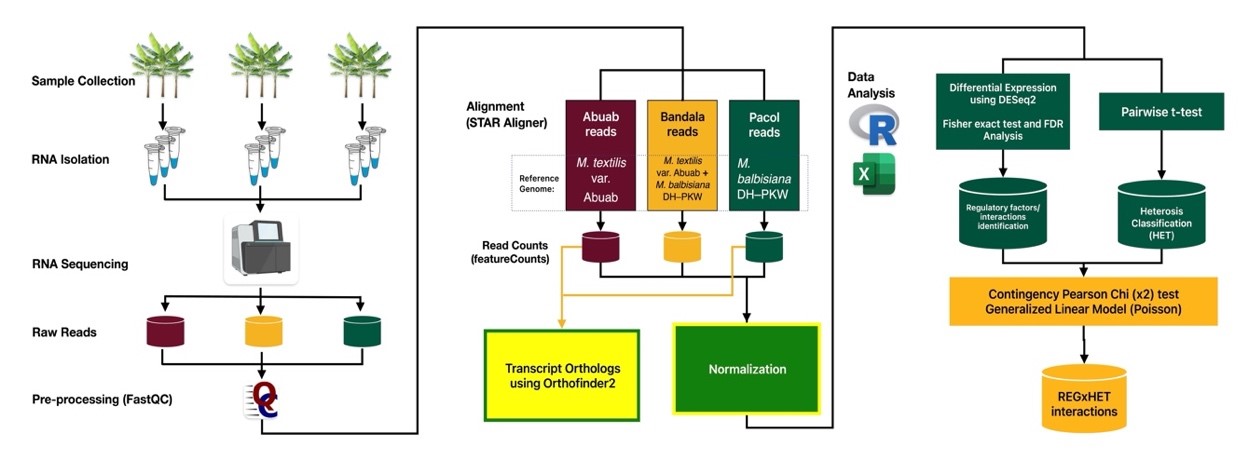

### Supp. Fig. 2

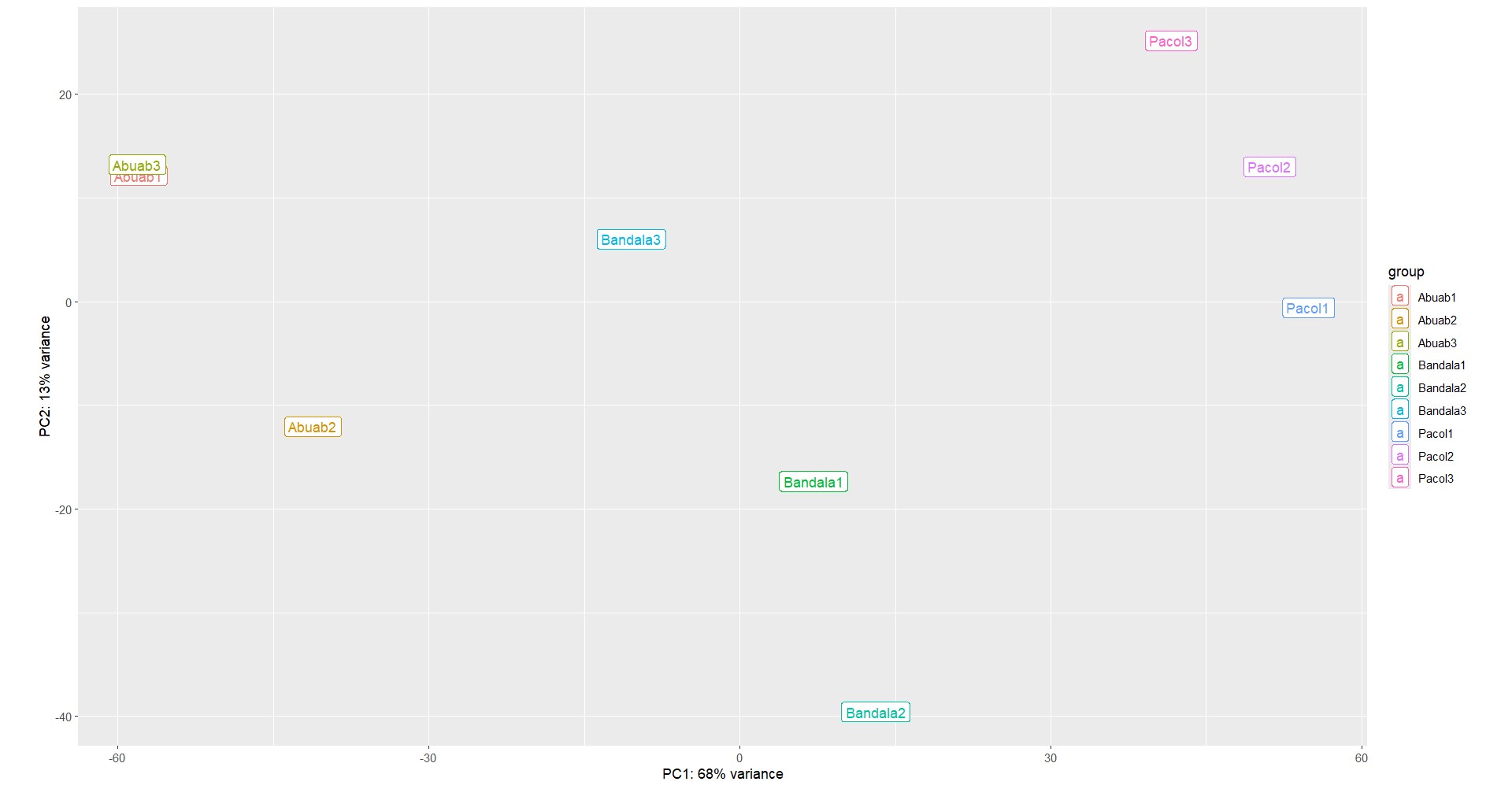

### Supp. Fig. 3

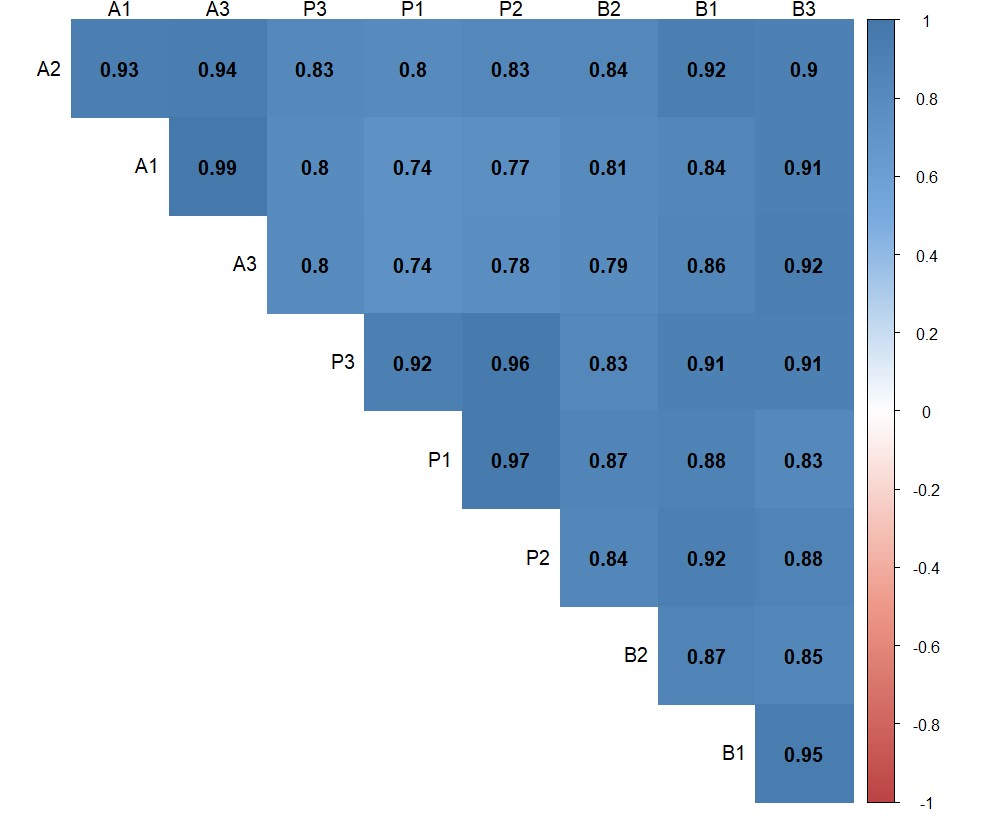

### Supp. Fig. 4

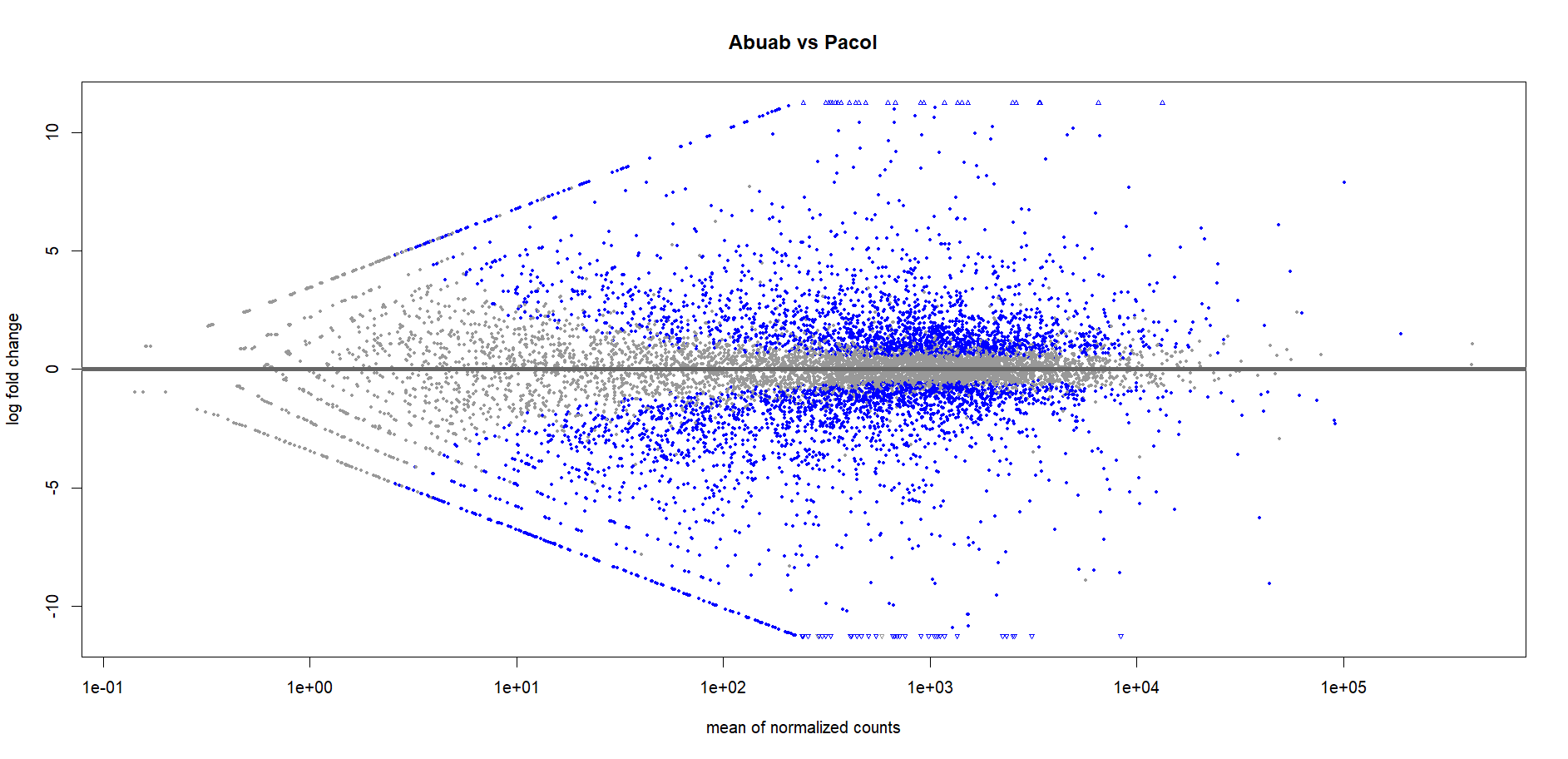

### Supp. Fig. 5

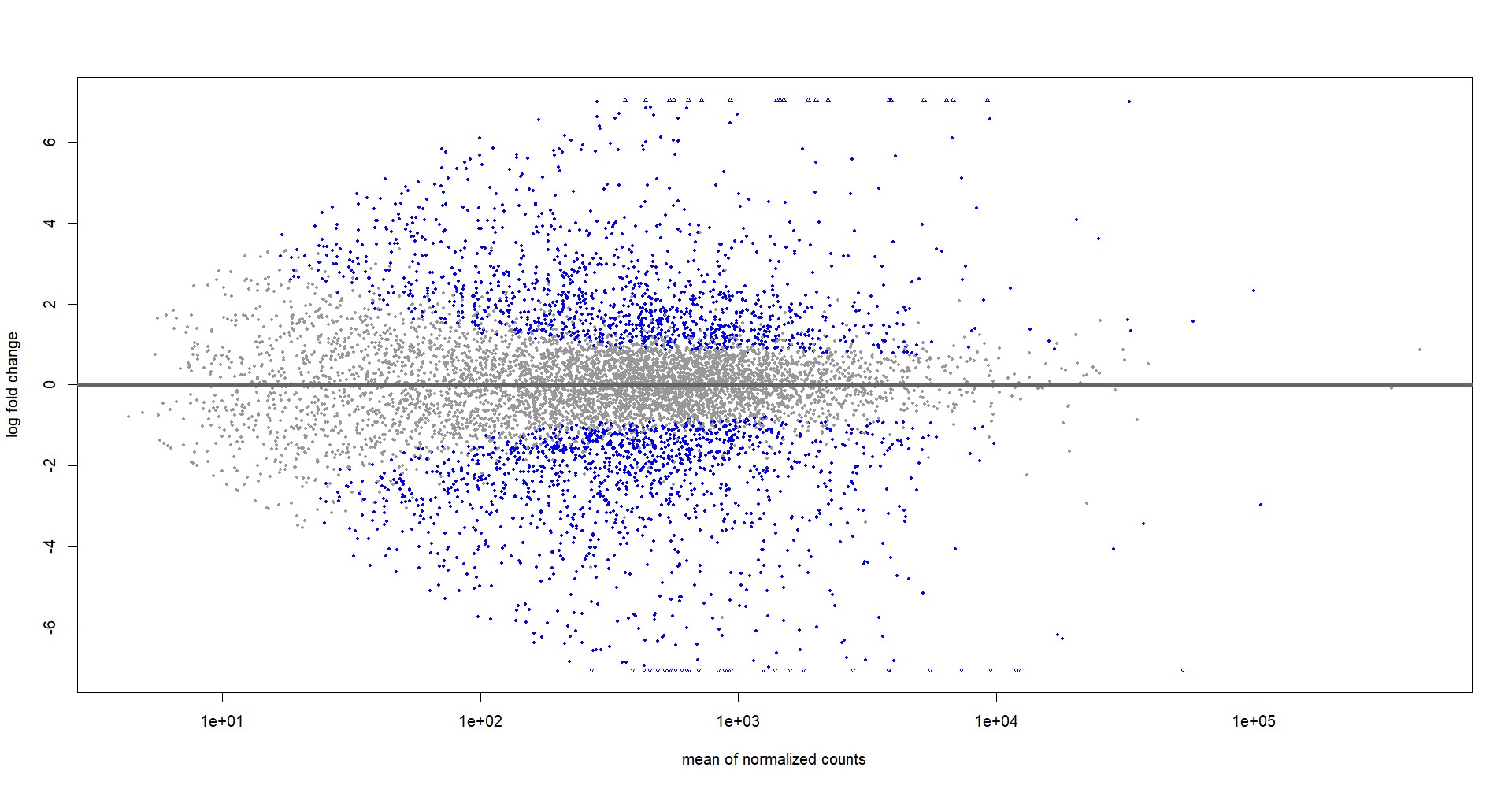

### Supp. Fig. 6

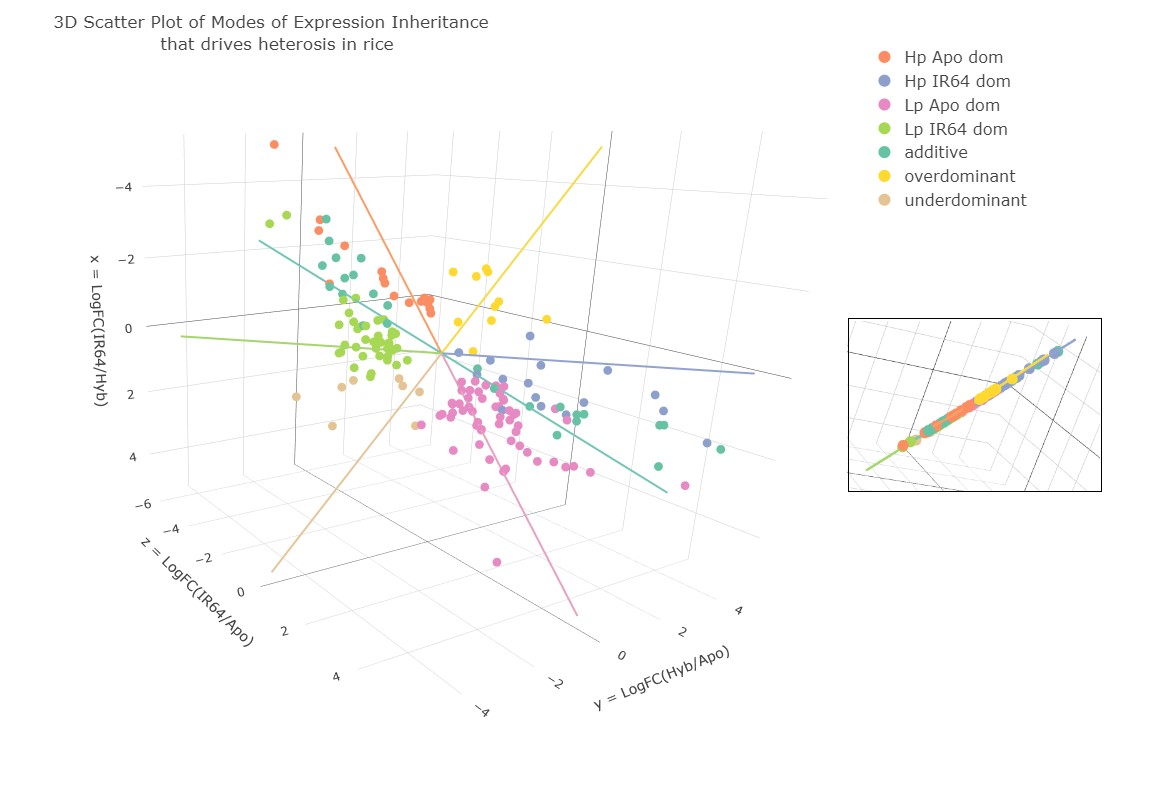

### Supp. Fig. 7

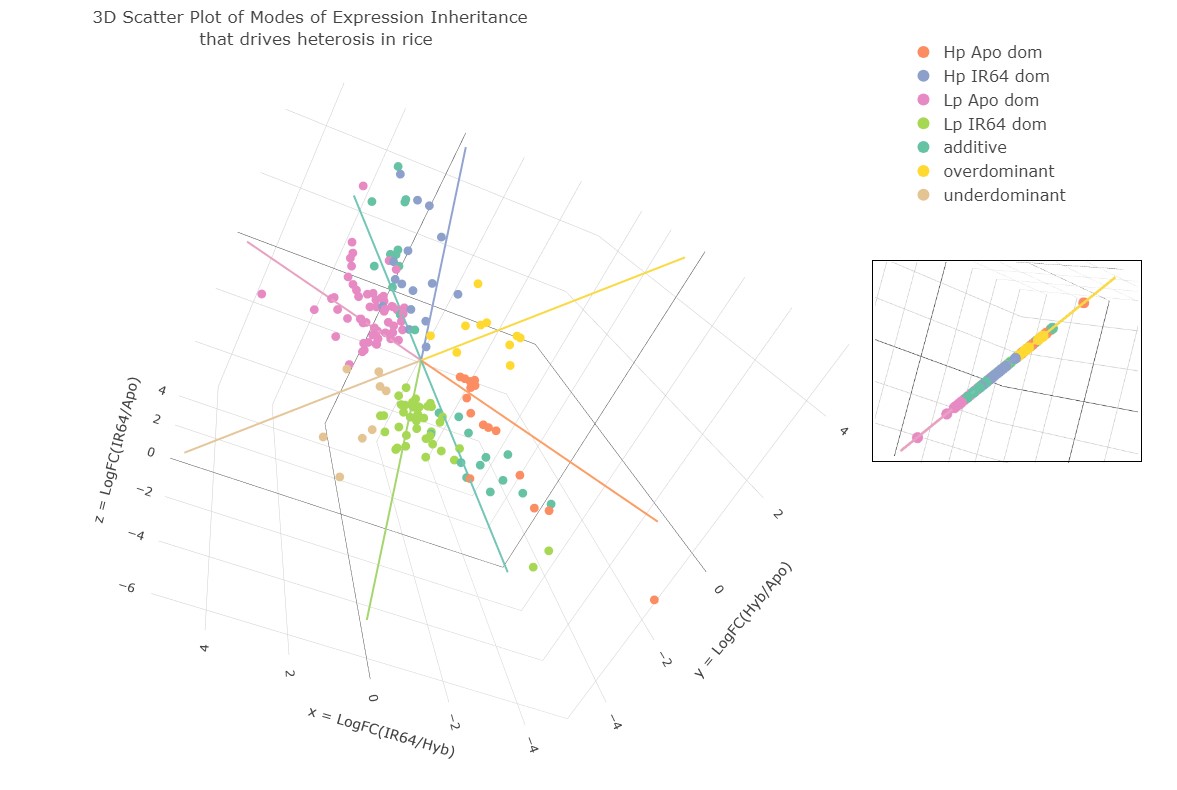

### Supp. Fig. 8

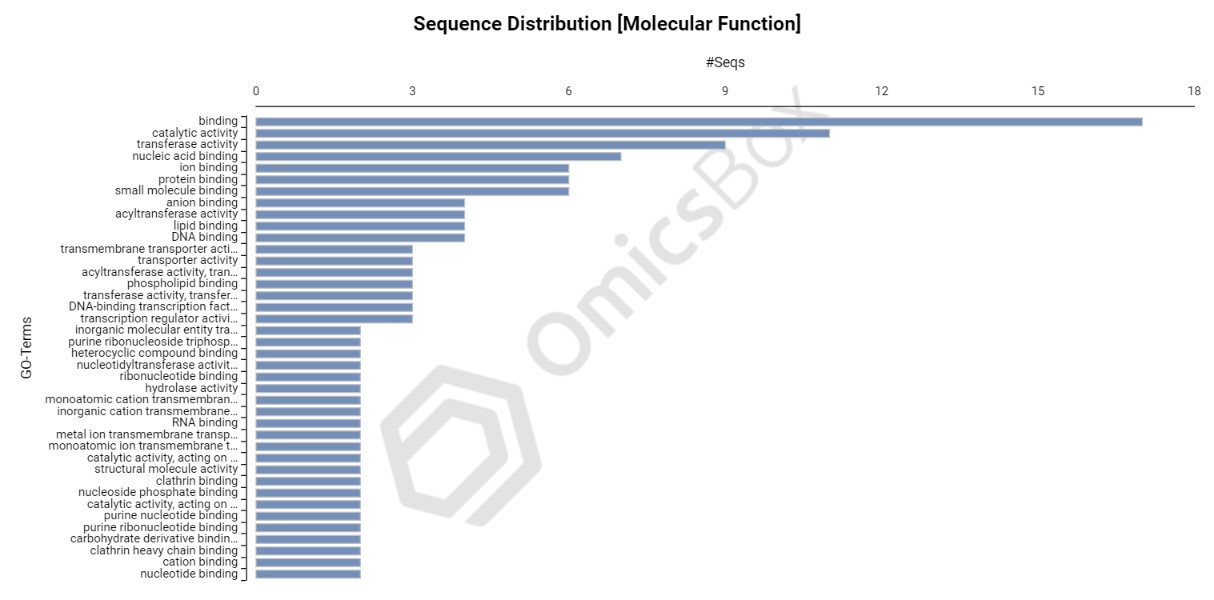

### Supp. Fig. 9

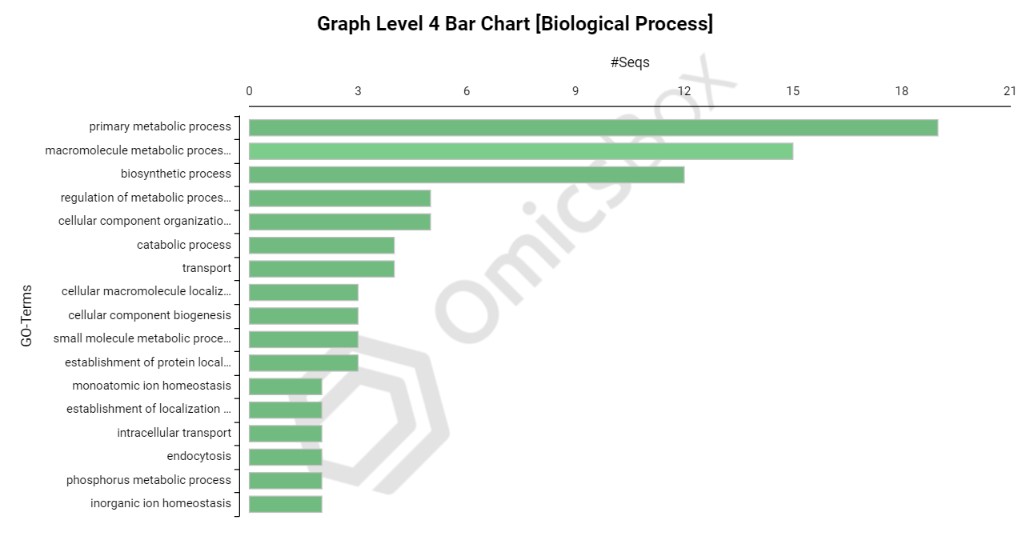
